# SPORTS1.0: A Tool for Annotating and Profiling Non-coding RNAs Optimized for rRNA- and tRNA-derived Small RNAs

**DOI:** 10.1101/296970

**Authors:** Junchao Shi, Eun-A Ko, Kenton M. Sanders, Qi Chen, Tong Zhou

## Abstract

High-throughput RNA-seq has revolutionized the process of small RNA (sRNA) discovery, leading to a rapid expansion of sRNA categories. In addition to the previously well-characterized sRNAs such as microRNAs (miRNAs), Piwi-interacting RNA (piRNAs), and small nucleolar RNA (snoRNAs), recent emerging studies have spotlighted on tRNA-derived sRNAs (tsRNAs) and rRNA-derived sRNAs (rsRNAs) as new categories of sRNAs that bear versatile functions. Since existing software and pipelines for sRNA annotation are mostly focused on analyzing miRNAs or piRNAs, here we developed the <u>s</u>RNA annotation <u>p</u>ipeline <u>o</u>ptimized for <u>r</u>RNA- and <u>t</u>RNA- derived <u>s</u>RNAs (SPORTS1.0). SPORTS1.0 is optimized for analyzing tsRNAs and rsRNAs from sRNA-seq data, in addition to its capacity to annotate canonical sRNAs such as miRNAs and piRNAs. Moreover, SPORTS1.0 can predict potential RNA modification sites based on nucleotide mismatches within sRNAs. SPORTS1.0 is precompiled to annotate sRNAs for a wide range of 68 species across bacteria, yeast, plant, and animal kingdoms, while additional species for analyses could be readily expanded upon end users’ input. For demonstration, by analyzing sRNA datasets using SPORTS1.0, we reveal that distinct signatures are present in tsRNAs and rsRNAs from different mouse cell types. We also find that compared to other sRNA species, tsRNAs bear the highest mismatch rate which is consistent with their highly modified nature. SPORTS1.0 is an open-source software and can be publically accessed at https://github.com/junchaoshi/sports1.0.

## Introduction

Expanding classes of small RNAs (sRNAs) have emerged as key regulators of gene expression, genome stability, and epigenetic regulation [1,2]. In addition to the previously well-characterized sRNA classes such as microRNAs (miRNAs), Piwi-interacting RNA (piRNAs), and small nucleolar RNA (snoRNAs), recent analysis of sRNA-seq data has led to the identification of expanding novel sRNA families. These include tRNA-derived sRNAs (tsRNAs; also known as tRNA-derived fragments, tRFs) and rRNA-derived sRNAs (rsRNAs) [3]. tsRNAs and rsRNAs have been discovered in a wide range of species with evolutionary conservation, supposedly due, in part, to the highly conservative sequence of their respective precursors, *i.e*., tRNAs and rRNAs [3]. Interestingly, tsRNAs and rsRNAs have been abundantly found in unicellular organisms (*e.g*., protozoa), where canonical sRNA pathways such as miRNA, siRNA, and piRNAs are entirely lacking [4−6]. The dynamic regulation of tsRNAs and rsRNAs in these unicellular organisms suggests that they are among the most ancient classes of sRNAs for intra- and inter-cellular communications [7].

Moreover, recent emerging evidence from mammalian species have highlighted the diverse biological functions mediated by tsRNAs, including regulating ribosome biogenesis, translation initiation, retrotransposon control, cancer metastasis, stem cell differentiation, neurological diseases, and epigenetic inheritance [3,8−15]. Although tsRNAs are known to be involved in regulating these processes at both post-transcriptional and translational levels [11,14,16], the exact molecular mechanisms of how tsRNAs exert their functions have not been fully understood. Compared to tsRNAs, rsRNAs are more recently discovered and also exhibit tissue-specific distribution. Dynamic expression of rsRNAs is associated with diseases such as metabolic disorders and inflammation [17−19]. The diverse biological functions of tsRNAs and rsRNAs and their strong disease associations are now pushing the new frontier of sRNA research.

Currently, there are multiple existing general sRNA annotation software and pipelines [20−24], and some have been developed aiming to analyze tsRNAs [25−27]. However, there still lack the specialized tools that can simultaneously analyze both tsRNAs and rsRNAs in addition to other canonical sRNAs. Here, we provide SPORTS1.0, which can annotate and profile canonical sRNAs such as miRNAs and piRNAs, and is also optimized to analyze tsRNAs and rsRNAs from sRNA-seq data. In addition, SPORTS1.0 can help predict potential RNA modification sites based on nucleotide mismatches within sRNAs.

## Method

The source code of SPORTS1.0 is written in *Perl* and *R*. The whole package and installation instructions are available on Github (https://github.com/junchaoshi/sports1.0). SPORTS1.0 can apply to a wide-range of species and the annotation references of 68 species are precompiled for downloading (Table S1).

The workflow of SPORTS1.0 consists of four main steps, *i.e*., pre-processing, mapping, annotation output, and annotation summary (**Figure 1**). SRA, FASTQ, and FASTA are the acceptable formats for data input. By calling Cutadapt [28] and *Perl* scripts extracted from miRDeep2 [29], SPORTS1.0 outputs clean reads by removing sequence adapters and discarding sequences with length beyond the defined range, and those with bases other than ATUCG. The clean reads obtained in pre-processing step are sequentially mapped against reference genome, miRBase [30], rRNA database (collected from NCBI), GtRNAdb [31], piRNA database [32,33], Ensembl [34] and Rfam [35], upon users’ setting. sRNA sequences are first annotated by Bowtie [36]. Next, a *Perl* script precompiled in SPORTS1.0 is used to identify the locations of tsRNAs regarding whether they are derived from 5’ terminus, 3’ terminus, or 3’CCA end of tRNAs. Then an *R* script precompiled in SPORTS1.0 is applied to obtain rsRNA expression level and positional mapping information regarding their respective rRNA precursors (5.8S, 18S, 28S, *etc*.).

**Figure 1.**
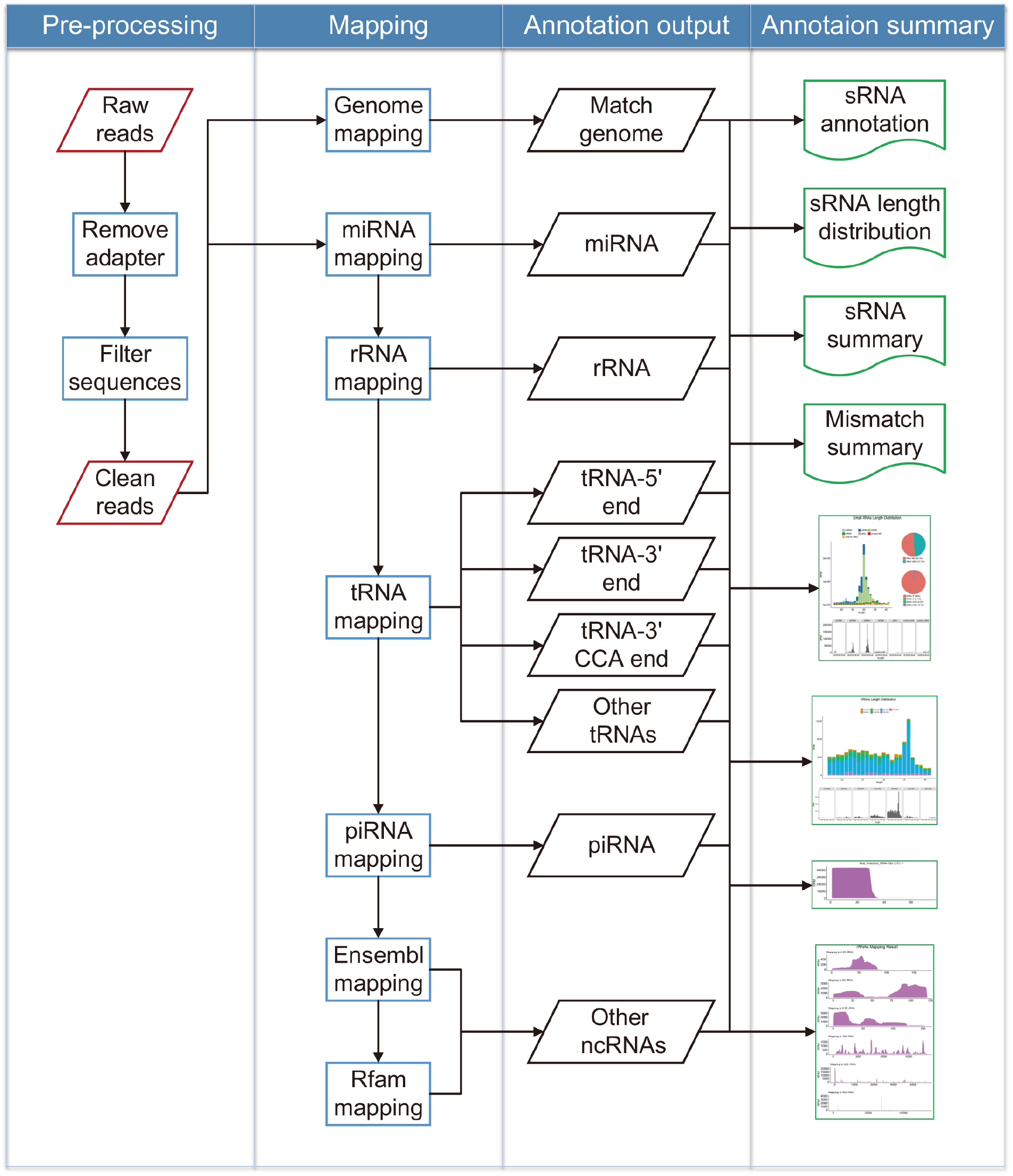
Workflow of SPORTS1.0. SPORTS1.0 contains four main steps, *i.e*., pre-processing, mapping, annotation output, and annotation summary, as outlined in the figure.

SPORTS1.0 can also be used to analyze sequence mismatch information if mismatches are allowed during alignment process. Such information can help predict potential modification sites that have caused nucleotide misincorporation during the reverse transcription (RT) process as previously reported [37]. In the current version, a mismatch site is designated using criteria as previously described [37]. Binomial distribution is used to address whether the observed mismatch enrichment is significantly higher than the base-calling error. Here, we define *p_err_* as the base-calling error rate, *n_ref_* as the number of nucleotides perfectly fitted to the reference sites, *n_mut_* as the number of mismatched nucleotides, and *n_tot_* as the sum of *n_ref_* and *n_mut_*. The probability of observing not larger than *k* perfectly matched nucleotides out of *n_tot_* can be calculated as:

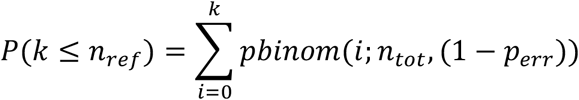

SPORTS1.0 provides two methods to evaluate *n_mut_* number. The first option is to simply calculate *n_mut_* as the read number of sequences containing particular mismatches. Since some sequences may align to multiple reference loci, using this method may result in an increased false-positive rate. A second method is thus included, in which read number of sequences from multiple matching loci are uniformly distributed (based on the assumption that each of these multiple sites will equally express RNAs) and consequently generates an adjusted *n_mut_*.

SPORTS1.0 summary output includes annotation details for each sequence and length distribution along with other statistics. (See sample output **Figure 2** and **Figure 3**, Table S2 and Table S3). User guideline is provided online (https://github.com/junchaoshi/sports1.0).

## Results

As an example, we used SPORTS1.0 to analyze sRNA-seq datasets from mouse sperm (GSM2304822 [38]), bone marrow cells (GSM1604100 [39]), and intestinal epithelial cells (GSM1975854 [40]) samples. Graphic output by SPORTS1.0 reveals distinct sRNA profiles in sperm (**Figure 2A**), bone marrow cells (Figure 2B), and intestinal epithelial cells (Figure 2C) samples. tsRNAs and rsRNAs are found equally or more abundantly than previously well-known miRNAs or piRNAs (length distribution data for each type of sRNA are exemplified in Table S2). In particularly, tsRNAs are dominant in sperm, rsRNAs are highest in bone marrow cells, and intestinal epithelial cells contains an appreciable amount of both tsRNAs and rsRNAs in addition to a miRNA peak.

**Figure 2.**
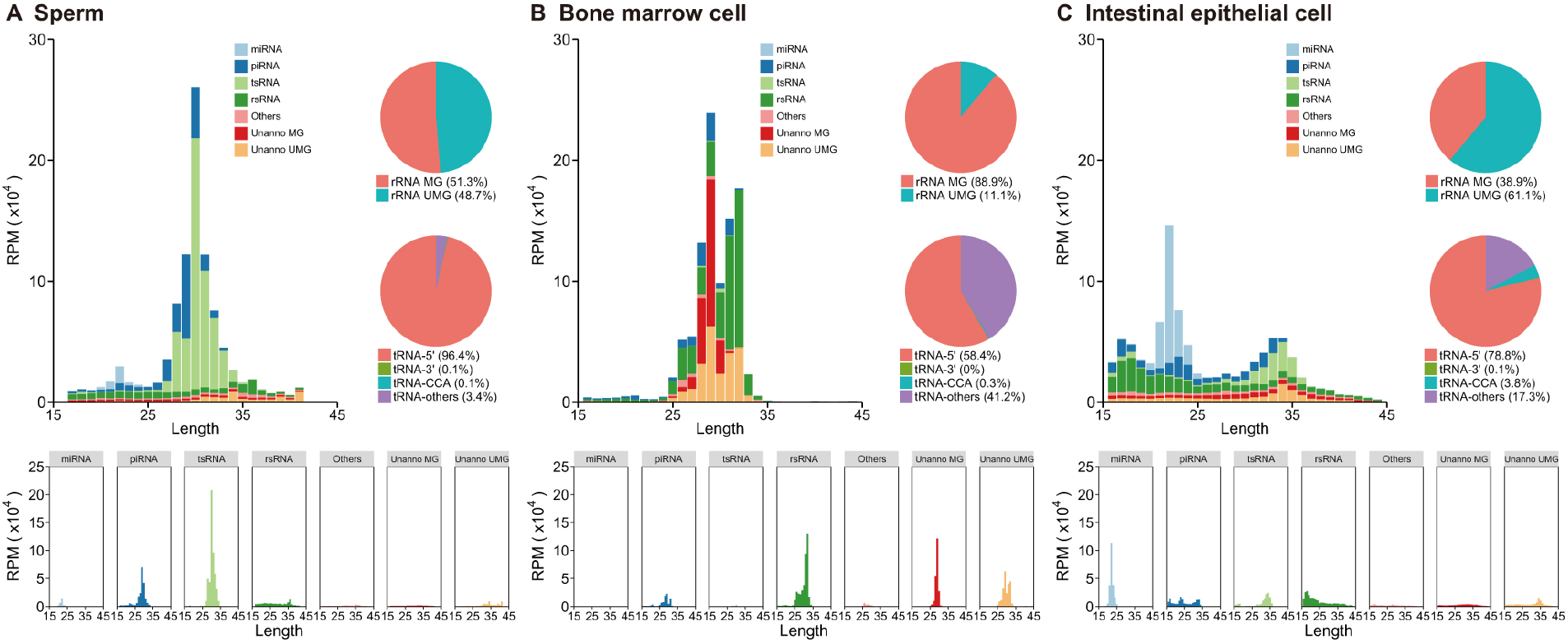
Exemplary annotation and profiling of sRNA-seq datasets generated by SPORTS1.0. Categorization and length distribution analysis of different sRNA types in mouse sperm (**A**), bone marrow cell (**B**), and intestinal epithelial cell (**C**) samples. RPM, reads per million clean reads; Unanno: unannotated; MG: match genome; UMG: unmatch genome.

Importantly, SPORTS1.0 found an appreciable portion of rsRNAs annotated in sperm (48.7%), bone marrow cell (11.1%) and intestinal epithelial cell (61.1%) samples that are previously deemed as “unmatch genome” (UMG) (Figure 2A-C upper pie-chart). This is because these newly annotated rsRNAs are derived from rRNA genes (rDNA), which were not assembled and shown in current mouse genome (mm10) [41], and thus were discarded before analysis by previous sRNA analyzing pipelines. SPORTS1.0 can now annotate and analyze these rsRNAs, including providing the subtypes of rRNA precursors (5.8S, 18S, 28S, *etc*.) from which they are derived from (**Figure 3A-C**), as well as the loci mapping information (Figure 3D−F). Interestingly, our analyses revealed that the specific loci that generate rsRNAs are completely distinct among sperm, bone marrow cell, and intestinal epithelial cell samples (Figure 3D-F), suggesting distinct biogenesis and functions of these rsRNAs. Similarly, SPORTS1.0 also revealed tissue-specific landscape of tsRNAs in terms of their relative abundance (Figure 2A-C lower pie chart) and the tRNA loci where they are derived from (5’ terminus, 3’ terminus, 3’CCA end, *etc*.) (**Figure 4**, and Figure S1−3). Since tsRNAs from different loci bear distinct biological functions [3], the tissue-specific tsRNA composition may represent features that define the unique functions of respective tissue/cell types.

**Figure 3.**
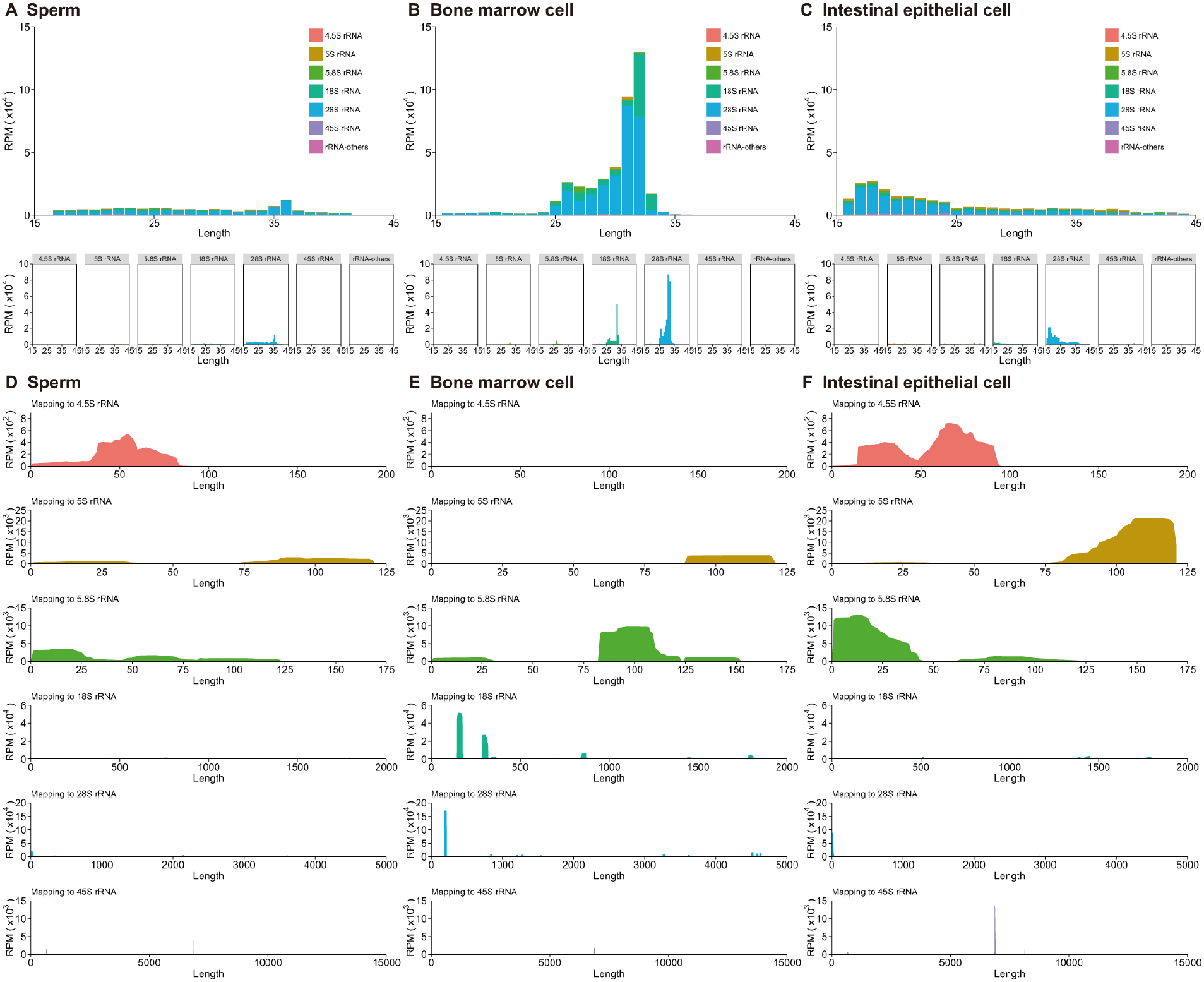
Cell-specific rsRNA profiles revealed by SPORTS1.0. Subtypes of rRNA precursors (5.8S, 18S, 28S, *etc*.) for rsRNAs from mouse sperm (**A**), bone marrow cell (**B**), and intestinal epithelial cell (**C**) samples. Comparison of rsRNA-generating loci from different rRNA precursors reveals distinct pattern between sperm (**D**), bone marrow cell (**E**), and intestinal epithelial cell (**F**). RPM, reads per million clean reads.

**Figure 4.**
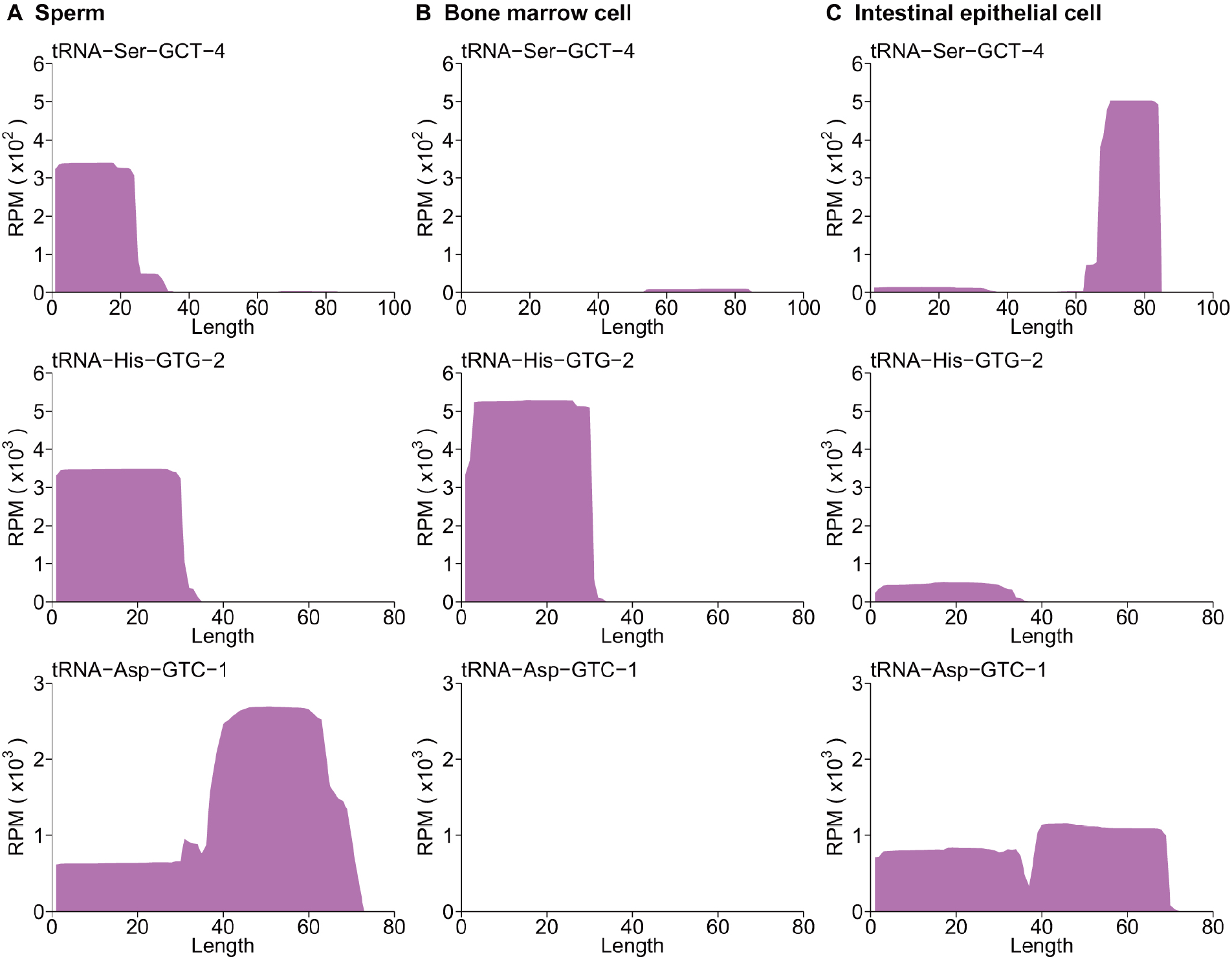
Cell-specific tsRNA profiles revealed by SPORTS1.0. Examples of 3 cell-specific tsRNA profiles revealed in mouse sperm (**A**), bone marrow cell (**B**), and intestinal epithelial cell (**C**) samples. Full tsRNA mapping results against tRNA loci are included in Figure S1−S3 for sperm (Figure S1), bone marrow cell (Figure S2), and intestinal epithelial cell (Figure S3) respectively. RPM, reads per million clean reads.

In addition, SPORTS1.0 also revealed distinct mismatch rates among different types of sRNAs (**Figure 5** and Table S3), with tsRNAs showing the highest. The detected mismatch sites represent the modified nucleotides that might have caused misincorporation of nucleotides during the RT process. The relatively higher mismatch rate detected in tsRNA sequences is consistent with their highly modified nature. The mismatch sites detected by SPORTS1.0 could provide a potential source for further analyses of RNA modifications within sRNAs.

**Figure 5.**
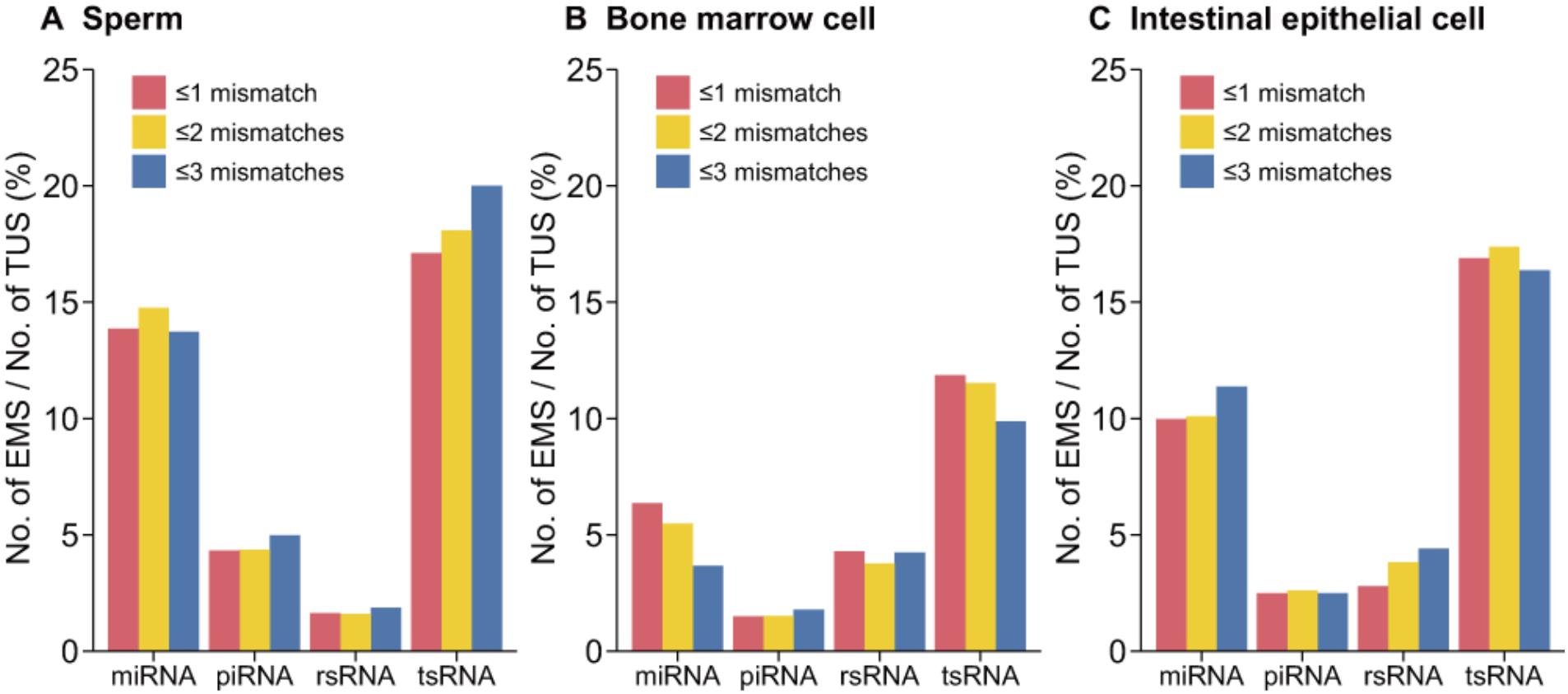
sRNA mismatch statistics by SPORTS1.0. The percentage of unique sequences that contain significantly-enriched mismatches out of total number of unique sequences from each subtype of sRNAs (miRNAs, piRNAs, tsRNAs, and rsRNAs) is provided for sperm (**A**), bone marrow cell (**B**), and intestinal epithelial cell (**C**) samples. EMS: enrichment mismatch sequences; TUS: total unique sequences.

Finally, SPORTS1.0 can analyze sRNAs of a wide range of species, depending on the availability of their reference genome and sRNA sequences (**Figure 6** and Table S**1**). The species to be analyzed and their associated sRNA references are subject to update in future versions, or can be customized by the end users.

**Figure 6.**
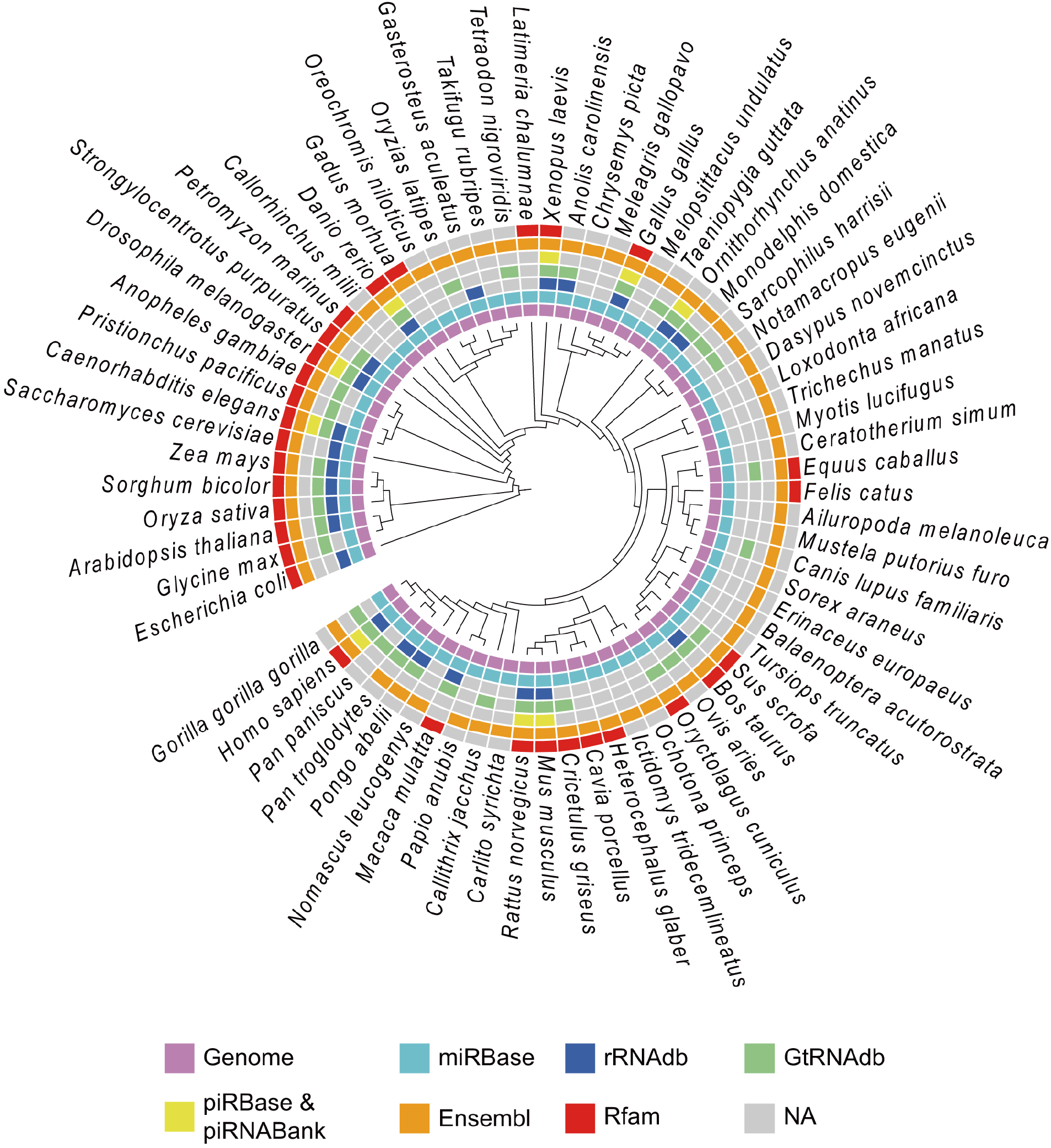
Species recompiled for analysis by SPORTS1.0. The 68 species and their respective reference database included in SPORTS1.0 precompiled for analysis.

## Conclusion

SPORTS1.0 is an easy-to-use and flexible pipeline for analyzing sRNA-seq data across a wide-range of species. Using mice as example, SPORTS1.0 provides a far more complicated sRNA landscape than having been previously seen, highlighting a tissue-specific dynamic regulation of tsRNAs and rsRNAs. SPORTS1.0 can also predict potential RNA modification sites based on nucleotide mismatches within sRNAs, and show a distinct pattern between different sRNA types. SPORTS1.0 may set the platform for many future new discoveries in biomedical and evolution research that is related to sRNAs.

> The real voyage of discovery consists not in seeking new landscapes, but in looking with new eyes.
>
> — -Marcel Proust

## Authors’ contributions

JS, TZ, and QC conceived the idea and wrote the manuscript. JS and TZ developed the SPORTS1.0 software and analyzed the RNA-seq data. JS, EK, KMS, QC, and TZ contributed to the interpretation of the results. All authors read and approved the final manuscript.

## Acknowledgments

We want to thank Songjia Fan and Tin Nguyen for their constructive suggestions for the manuscript. This work is supported by Start-up funds for Zhou and Chen labs from University of Nevada, Reno School of Medicine, and from National Institutes of Health, USA (Grant Nos. R01DK091336 and P01DK041315 to KMS; Grant Nos. R01HD092431 and P30GM110767-03 to QC).

## Supplementary materials

**Figure S1 The mouse sperm tsRNA mapping results against tRNA loci revealed by SPORTS1.0** Mapping result for each annotated tsRNA was provided.

**Figure S2 The mouse bone marrow cell tsRNA mapping results against tRNA loci revealed by SPORTS1.0** Mapping result for each annotated tsRNA was provided.

**Figure S3 The mouse** intestinal epithelial cell **tsRNA mapping results against tRNA loci revealed by SPORTS1.0** Mapping result for each annotated tsRNA was provided.

**Table S1 Table S1 The list of 68 species and their respective reference database that are precompiled in SPORTS1.0 ready for analyses**

**Table S2 Table S2 Example output of SPORTS1.0 which includes annotation for each sequence (A), length distribution information (B) and expression level of each annotated category (C) for dataset GSM2304822**

**Table S3 Table S3 Example output of SPORTS1.0 for sRNA sequence mismatch analysis for dataset GSM2304822 under the alignment criteria of mismatch ≤ 1 (A), ≤ 2 (B), and ≤ 3 (C)**

